# Transcripts of vaccinia virus postreplicative genes do not contain a 5’ methylguanosine cap

**DOI:** 10.1101/2020.07.15.204867

**Authors:** Václav Vopálenský, Michal Sýkora, Zora Mělková, Tomáš Mašek, Martin Pospíšek

## Abstract

Vaccinia virus (VACV) is a prototypical poxvirus originally used for eradication of smallpox. Investigation into VACV mRNAs carried out almost half a century ago substantially contributed to the fundamental discovery of the 5’ mRNA cap, a hallmark of all eukaryotic and many viral mRNAs. VACV research also facilitated the identification and understanding of the general mechanism of 5’ mRNA cap synthesis. We characterized the VACV transcripts at the individual mRNA molecule level and found that vaccinia postreplicative mRNAs, containing nontemplated 5’ poly(A) leaders, surprisingly lack the 5’ cap structure *in vivo*. We also show that the lengths of the nontemplated leaders and the presence or absence of cap structures at the 5’ mRNA ends are controlled by the initiator sequence of the VACV postreplicative promoters.

**One Sentence Summary:** The promoter sequence determines the synthesis of the 5’ cap and poly(A) leaders in vaccinia virus postreplicative mRNAs.

## Main text

Poxviruses belong to a family of large cytoplasmic animal and arthropod double-stranded DNA viruses, including two viruses that exclusively infect humans, namely, molluscum contagiosum virus and variola virus, the latter being the causative agent of smallpox. Before its eradication, variola virus killed more humans than all other infectious diseases collectively (*1*). Some poxviruses possess the ability to cross the interspecies barrier and can be transmitted from animal hosts to humans. From this perspective, the most effective poxvirus is the monkeypox virus, for which direct human-to-human transmission has also been reported. The numbers of human infections caused by monkeypox have been increasing in Central and West Africa since the end of the smallpox vaccination, and several imported outbreaks were also reported in the United States of America, the United Kingdom, Israel and Singapore in recent years (*2, 3*). Current advances in synthetic biology and genetic engineering have led to the recent reassessment of smallpox among the most dangerous bioterrorism agents in category A, where it newly received the highest risk-priority score together with anthrax (*4*). The characteristics of animal and human poxviruses, global travel and decreased global political stability have thus led to the inclusion of poxviruses among potential emerging agents of the next deadly pandemics for which direct human-to-human transmission has also been reported. Therefore, programs aimed at developing better smallpox vaccines and treatment have been recently been revitalized worldwide (*5*), and new data about the replication of poxviruses and their interaction with host cells and the immune system are needed.

Gene expression of vaccinia virus (VACV), a prototypical member of the *Orthopoxvirus* family used in the smallpox eradication campaign (*6*), proceeds as a tightly regulated cascade of sequential transcription of early, intermediate and late genes (gene time classes, GTCs) and is regulated primarily at the transcription initiation level (*7*). VACV-encoded RNA polymerase is a multisubunit enzyme (*8*) that exists in two distinct forms that are specific for transcription of early and postreplicative genes (*9*). The 5’ ends of early and postreplicative VACV transcripts are traditionally considered to be protected by the 7-methygluanosine cap structure (m^7^G) (*10, 11*), which is synthesized by the VACV mRNA capping enzyme (*12, 13*). The promoter composition and transcription regulation of VACV intermediate and late genes differ from those of early genes. Mutational analyses and transcriptome sequencing identified an essential 15-nt-long A/T-rich promoter core sequence of the VACV early genes located ~12 nt upstream of the transcription start site (TSS) (*14*). Promoters of VACV intermediate and late genes contain 8–11-nt A/T-rich core elements located ~11 and 6 nt upstream of the initiator region (INR), respectively (*15–17*). Using VACV transcriptome sequencing, consensus promoter sequences of intermediate and late genes have been determined (*14, 18*). Postreplicative mRNAs have relatively short 5’ untranslated regions (*18*), and their initiation nucleotide is adenosine both *in vivo* and *in vitro* (*11, 19*). Mapping of the 5’ ends of the VACV intermediate and late mRNAs by primer extension revealed 5’ poly(A) leaders approximately 35 nt in length that were not encoded by the viral genome (*20–23*). These nontemplated 5’ poly(A) leaders are likely produced by repeated RNA polymerase sliding on the INR *in vivo* and *in vitro* (*17, 24*). Transcription initiation sites for the vast majority of VACV postreplicative genes are located within 25 nt of the translation initiation codon (*18*). The exact biological role of the 5’ poly(A) leaders in postreplicative transcripts has not been known for long, but it has been shown recently that the 5’ poly(A) leaders provide viral mRNAs a translational advantage over the host cell transcripts in infected cells (*25*). We recently showed that VACV RNA polymerase belongs to the monophyletic group with RNA polymerases encoded by yeast virus-like elements (VLEs), and the VLE promoters are related to those of poxviruses (*26*). VLE mRNAs also contain 5’ poly(A) leaders of variable length that are not complementary to the VLE DNA (*27*). We provided evidence that 5’ poly(A) leaders of VLE mRNAs are formed by a mechanism similar to the synthesis of 5’ poly(A) leaders of the VACV postreplicative transcripts (*26*). We further demonstrated that 5’ highly polyadenylated VLE mRNAs do not contain the m^7^G cap moiety at their termini. Similarities between VACV and VLEs in terms of RNA polymerases, promoters and 5’ mRNA poly(A) leaders (*26, 27*) prompted us to perform a detailed characterization of VACV mRNAs at the level of individual mRNA molecules.

The presence of the 5’ cap at the 5’-ends of VACV transcripts is commonly recognized, and VACV was among the first DNA viruses in which the 5’ mRNA cap moiety was identified (*10, 28, 29*). By analysis of the total VACV early and postreplicative mRNAs synthesized *in vivo*, it has been estimated that the 5’ cap structure occurs once per thousand nucleotides in early transcripts and once per 1,250 nucleotides in postreplicative mRNA transcripts (*11*). However, the presence of the 5’ cap structure in individual VACV transcripts has never been tested. To achieve this goal, we decided to apply a modification of the rapid amplification of 5’ cDNA ends (5’ RACE) method that we successfully applied for analysis of 5’ polyadenylation and capping of VLE mRNAs (*26, 27*). We purified total RNA from HeLa cells infected and mock-infected with VACV and investigated the 5’ ends of selected mRNAs covering all three GTCs. Consistent with the published results, our analysis revealed that a vast majority of VACV early mRNAs, which are controlled by promoters lacking INR (represented here by the genes *I4L*, *J6R*, *H5R*), lacked 5’ poly(A) leaders and contained a 5’ cap. In contrast, only approximately half of the VACV early transcripts controlled by promoters containing the INR, and thus bearing 5’ poly(A) leaders, contained a 5’ cap structure (represented by the genes *A5R* and *D12L*) (Figs. 1A, S1; Table S2). VACV intermediate transcripts were capped even less frequently. We identified a 5’ cap moiety in 33% and 11% mRNAs of the intermediate genes *G8R* and *A1L*, respectively (Figs. 1B, S2; Table S2). Capped late VACV mRNAs were detected only sporadically (*C3L*) or not at all (*A17L*) (Figs. 1C, S3; Table S2). Consistent with previous findings (*30*), we confirmed that the length of the 5’ poly(A) leader in VACV transcripts significantly increases in each successive GTC (Fig. S6; Table S7). As in VLE mRNAs (*27*), the occurrence of the 5’ cap in VACV transcripts was significantly negatively correlated (r = −0.89, *p* < 0.01) with the increasing number of nontemplated adenosines in the 5’ poly(A) mRNA leaders (Figs. 2, S5; Table S6). The finding that the vast majority of VACV postreplicative transcripts contain almost exclusively uncapped 5’ poly(A) leaders is surprising and unexpected. However, this result is consistent with experiments showing a low requirement of VACV late mRNAs for the functional cap-dependent translation initiation pathway (*25, 31*) and the dependence on virus-induced phosphorylation of receptor for activated C kinase (RACK1) (*32*). Correspondingly, VLE mRNAs containing 5’ poly(A) leaders are loaded on polysomes independently of the cap-binding eIF4E translation initiation factor (*27*).

**Fig. 1.**
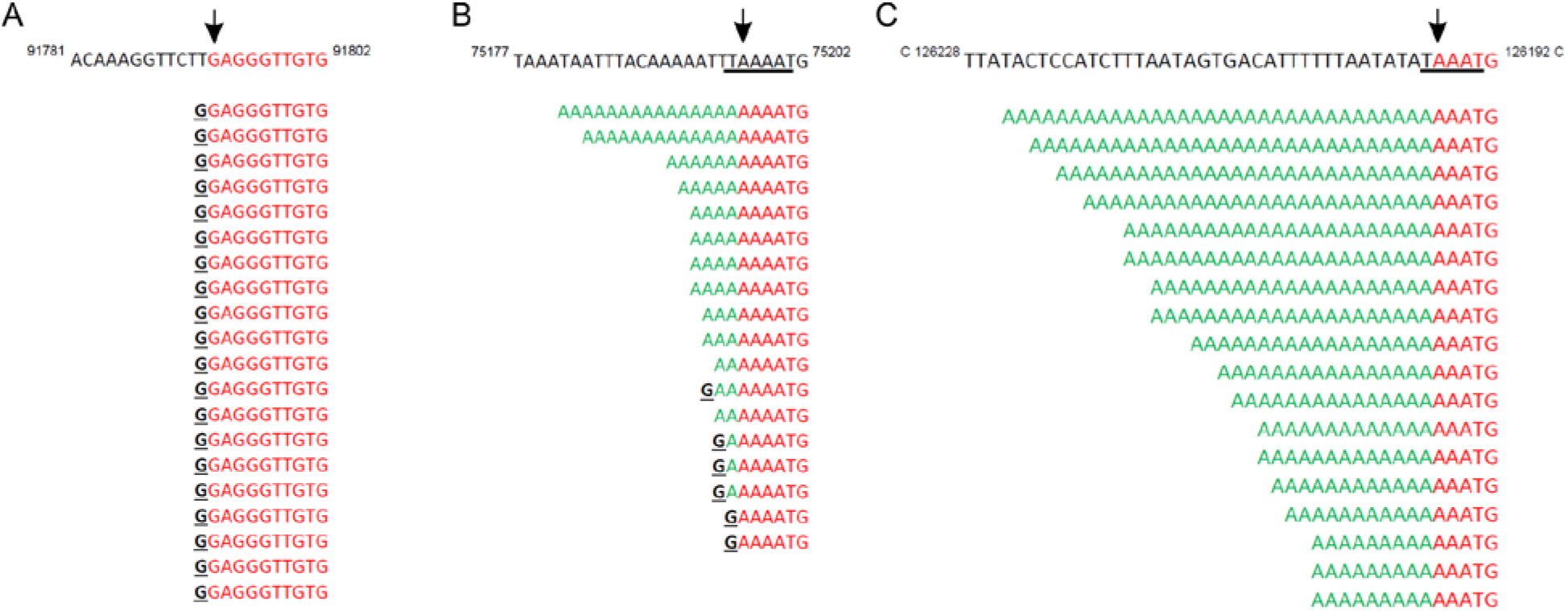
5’ RACE analysis of VACV transcripts. **(A)** *H5R* early gene transcripts are all capped (see also Fig. S1); the first 10 nucleotides of the 5’ untranslated region are displayed; the ATG translation initiation start codon is located 78 nt downstream from the TSS (Fig. S1). **(B)** Only a part of the *G8R* intermediate transcript contains 5’ cap moieties (see also Fig. S2). **(C)** All *C3L* late transcripts contain long 5’ poly(A) leaders and are not capped (see also Fig. S3). The upper sequence in each panel corresponds to the viral template DNA. The TSS (black arrow) and INR (underlined) were annotated according to (*14, 30*). If our TSS annotation differs, it is marked by an orange arrow. The sequences depicted below the template DNA represent individual sequenced cDNA clones. The 5’ untranslated regions are shown up to the ATG translation start codon, except in panel A. Nucleotides identical to the viral template DNA are labeled in red. Nucleotides added in a nontemplated manner are labeled in green. The guanosine residues corresponding to the 5’ mRNA cap are marked in black. All sequences are shown in the 5’-to-3’ orientation, regardless of their transcriptional orientation in the VACV genome.

**Fig. 2.**
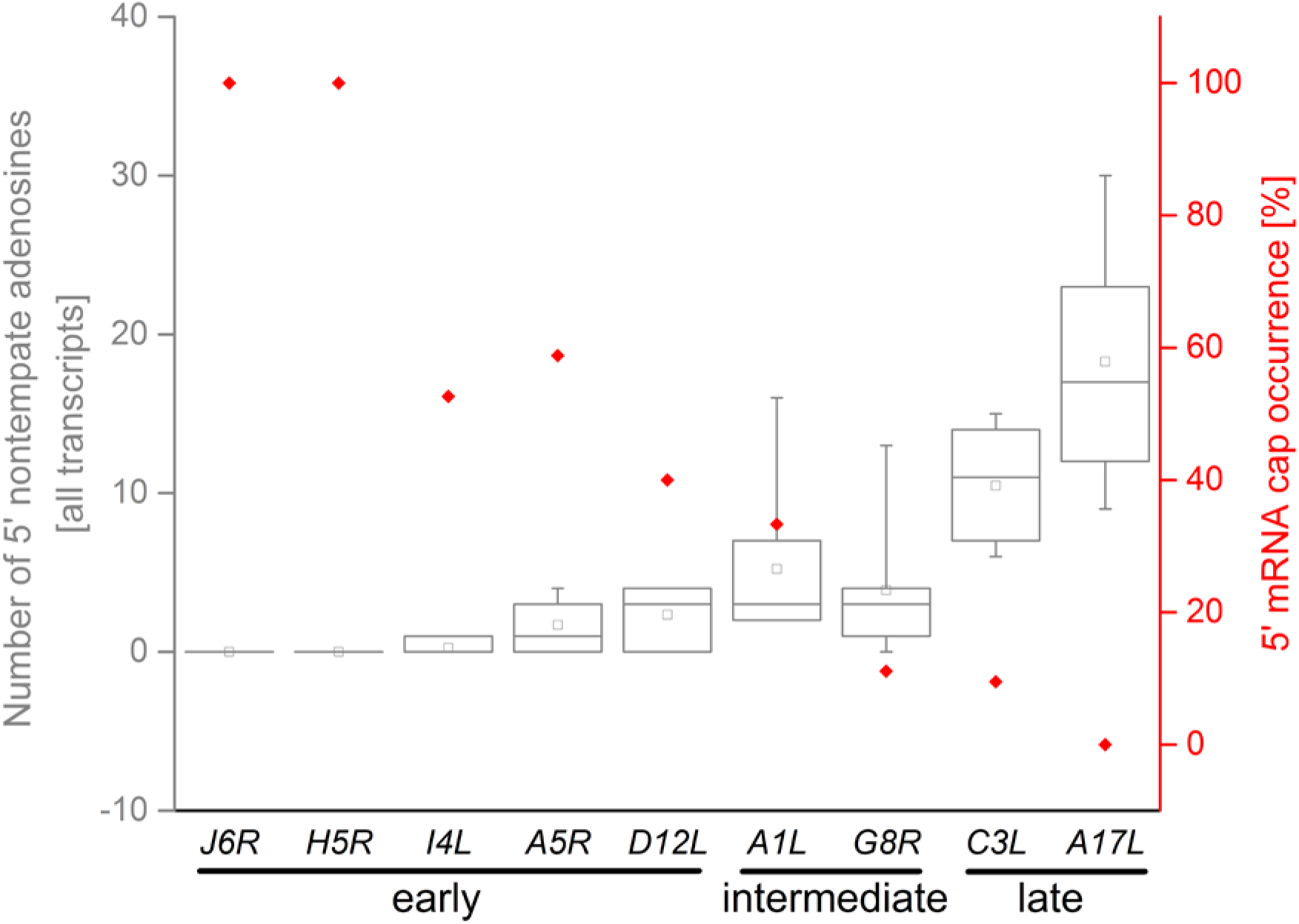
Negative correlation between the 5’ poly(A) leader length (left Y axis) and cap occurrence (right Y axis) in mRNAs from different successive GTCs represented by selected early, intermediate and late vaccinia genes (X axis). The box-whisker plot (in gray) represents the lengths of nontemplated 5’ poly(A) leaders. The lower bar, upper bar, bottom and top of the boxes represent the 10^th^ percentile, 90^th^ percentile and the first and third quartiles, respectively. The square dot and inner line in each box represent the mean and median, respectively. The proportions of 5’ capped transcripts [%] among all mRNAs tested for each particular gene are shown in red. In total, 166 individual cDNAs were used for this analysis (Figs. S1-S3, S5; Tables S2, S6).

The low occurrence of cap moieties at the 5’ ends of VACV late transcripts makes determination of the 5’ leader lengths of their capped variants technically challenging because it would require sequencing of unrealistically large numbers of 5’-RACE clones. Therefore, we decided to enrich 5’ capped transcripts of the selected VACV late genes by the RNA ligase-mediated amplification of cDNA ends (RLM-RACE) technique. RLM-RACE (*a.k.a*. oligo-capping) enables amplification of cDNA derived only from capped mRNAs by employment of three specific subsequent enzymatic reactions preceding the reverse transcription step. This procedure leads to a replacement of the original 5’ mRNA cap moiety with a short RNA oligonucleotide, which is then used after reverse transcription for subsequent PCR amplification (*33*). We applied the RLM-RACE approach for 5’ end mapping of VACV late genes, in which we expected the most striking difference in the lengths of capped and uncapped 5’ poly(A) leaders. Identical samples of total RNA purified from VACV-infected and mock-infected HeLa cells were used for the 5’-RACE and RLM-RACE techniques. Combined analyses of the 5’ RACE and RLM-RACE results clearly demonstrated that 5’ capped transcripts of both selected late VACV genes (*A17L* and *C3L*) tended to contain shorter 5’ poly(A) leaders than transcripts that lacked a 5’ mRNA cap moiety; this difference was statistically significant for the *C3L* mRNA at the *p*<0.01 level (Fig. 3; Table S3).

**Fig. 3.**
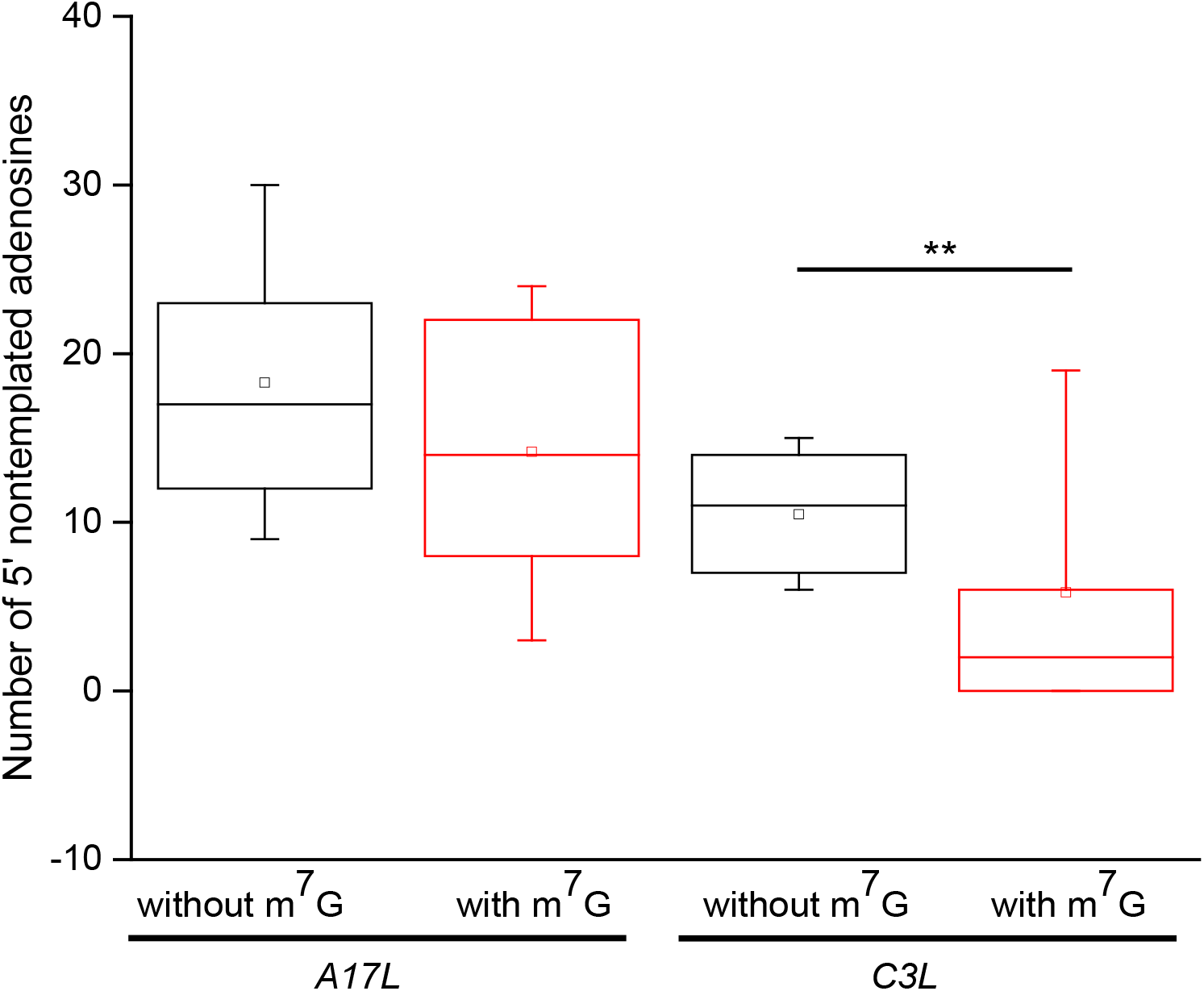
5’ capped late VACV mRNAs contain shorter 5’ poly(A) leaders than their uncapped counterparts. The box-whisker plot represents the lengths of nontemplated 5’ poly(A) leaders in *A17L* and *C3L* mRNAs that either contained (in red) or did not contain (in black) the 5’ cap moiety (m^7^G); the general description of the box plot is the same as that in Fig. 2. *p*<0.01 (**). In total, 96 sequences were used for the analysis (Fig. S4; Table S3).

Taking into account the negative correlation between the VACV 5’ mRNA poly(A) leader lengths and 5’ cap occurrence, we speculated that the 5’ polyadenylation rate of VACV mRNA directly affects m^7^G synthesis at the 5’ ends of VACV mRNAs. This possibility is further supported by our results showing that 5’ leaders of the capped late VACV transcripts were markedly shorter than 5’ leaders of the uncapped transcripts derived from the same genes (Fig. 3). These results might indicate the possibility that the length of the 5’ poly(A) mRNA leaders of VACV postreplicative transcripts calculated from whole-genome analyses of the VACV transcriptome might be underestimated, because only mRNAs containing the 5’ cap were used for deep sequencing (*30*). The medians of VACV intermediate transcript 5’ poly(A) leader lengths determined by Yang et al. and us are comparable and span 8 nt and 7–8 nt, respectively (*30*) (Fig. S2). However, the medians of VACV late transcript 5’ poly(A) leader lengths determined by cap-independent 5’ RACE were between 18–20 nt and as such were significantly longer than those determined by the cap-selective deep-sequencing method (11 nt, Fig. S3) (*30*). The latter readout is also in good agreement with the results obtained by us using the cap-selective RLM-RACE approach, yielding a median length of 8 nt for the VACV late mRNA 5’ poly(A) leaders (Fig. S4).

The INR oligo(A) stretch is commonly considered to be responsible for VACV RNA polymerase sliding and thus for 5’ leader synthesis. We wanted to test whether disruption of the continuous INR oligo(A) element can affect the length of the 5’ poly(A) leaders or the frequency of capping of VACV postreplicative mRNAs. We constructed reporter vectors bearing a gene encoding green fluorescent protein (EGFP), the expression of which was controlled by the viral *G8R* intermediate promoter containing either its wild-type INR sequence (TAAAAT; *pG8R^P^* - EGFP vector) or the point-mutated INR (TAA<u>C</u>AT; *pG8R^PM^*-EGFP vector). We transfected these reporter vectors into VACV-infected HeLa cells, purified total RNA 12 hours postinfection and subjected this RNA preparation to 5’ RACE analysis as described above. At least 19 independent cDNA clones corresponding to each wild-type and mutated INR variant were analyzed (Fig. 4A). Consistent with our expectation, the median lengths of the 5’ poly(A) leaders were comparable among mRNAs, the transcription of which was directed either from the viral genomic *G8R* promoter (3 nt) or its *G8R^P^* vector counterpart (2.5 nt; Fig. 4B; Table S4). In addition, the proportion of 5’ nontemplated poly(A) leader-containing mRNAs was comparable in both of these groups: 89.9% of viral *G8R* transcripts and 83.3% of *G8R^P^* vector mRNAs were 5’ oligoadenylated (Fig. 4C; Table S4). In contrast, a single point mutation in the *G8R* INR (*G8R^PM^*, TAA<u>C</u>AT) led to a significant decrease in the median and mean of the number of nontemplated adenosine nucleotides added to the 5’ ends of the reporter mRNAs (0 and 0.3, respectively; Fig. 4B, Table S4) and to a significant decrease in the fraction of 5’ polyadenylated mRNAs. The ratio of 5’ polyadenylated mRNAs to all *pG8R^PM^*-EGFP transcripts dropped to 31.6%. Consistent with the previously observed negative correlation between the length of 5’ poly(A) leaders and the occurrence of 5’ capping, reporter mRNAs controlled by point-mutated *G8R^PM^* INR were more frequently 5’ capped than reporter mRNAs, the transcription of which was directed by the wild-type promoter *G8R^P^* (57.9% vs 20.8%, respectively; Fig. 4D; Table S4). These results are also consistent with previously published data for 5’ polyadenylated VLE mRNAs (*27*).

**Fig. 4.**
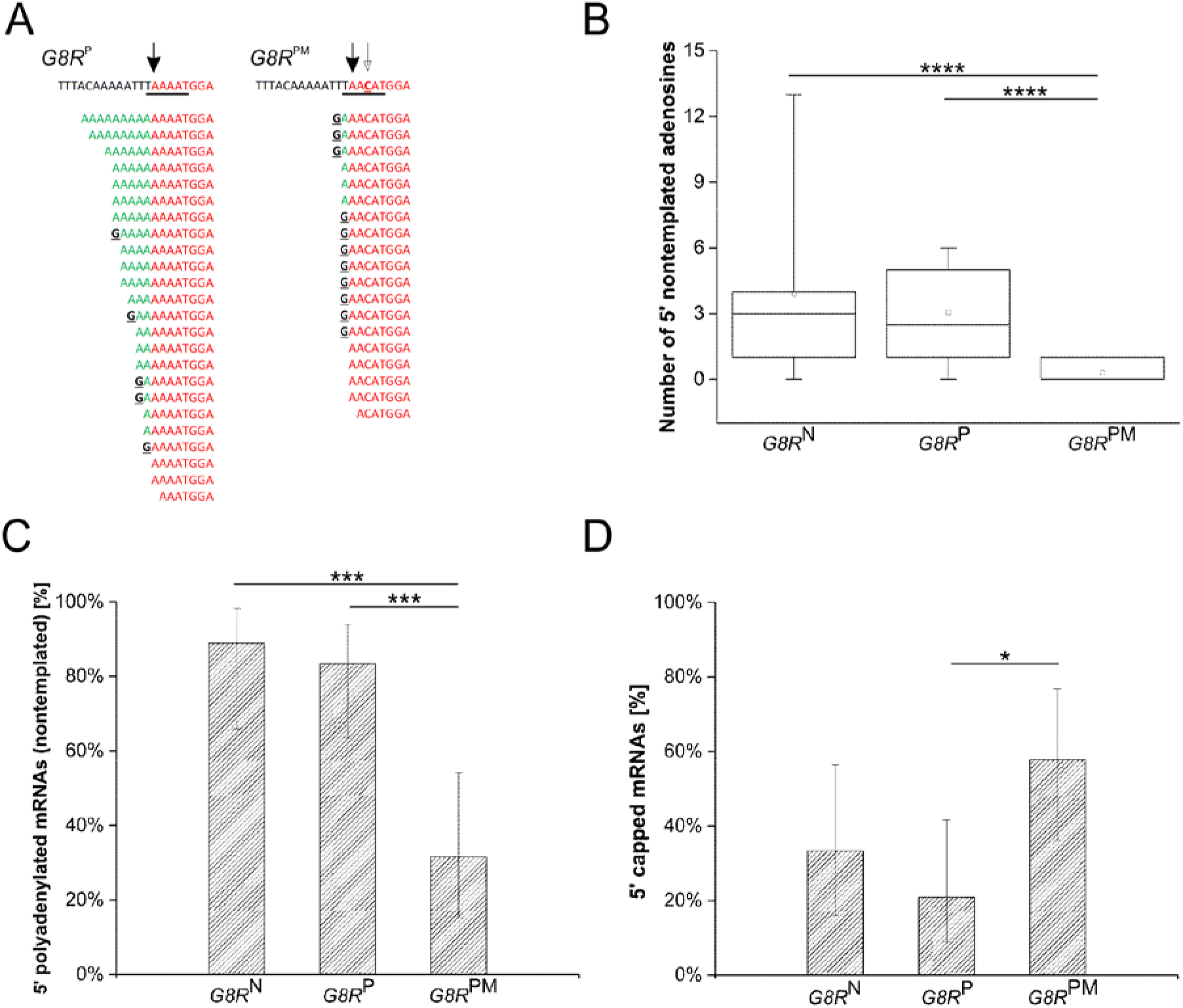
VACV INR controls 5’ end formation of viral mRNAs. **(A)** 5’ RACE analysis of EGFP reporter mRNAs transcribed from the intermediate wild-type *G8R* promoter (*G8R^P^*) or its point-mutated version (*G8R^PM^*). Single-nucleotide A/C substitution in *G8R* INR (empty arrow, underlined in red) resulted in the reduction in the number of added nontemplated adenosine residues and dramatic shortening of the 5’ poly(A) leaders. The upper sequence corresponds to the viral template DNA. TSS and INR are marked with a filled arrow and black underline, respectively. The sequences below represent individual sequenced cDNA clones. The 5’ untranslated regions are shown up to the ATG translation start codon. Nucleotides identical to the viral template DNA are labeled in red. Nucleotides added in a nontemplated manner are labeled in green. The guanosine residues corresponding to the 5’ mRNA cap are marked in black. **(B)** Statistical representation of panel A including results from 5’ RACE analysis of natural viral *G8R* mRNAs (*G8R^N^*; Fig. S2; Table S2) for comparison. The general description of the box plot is the same as that in Fig. 2. **** indicates *p*<0.0001 **(C)** Single A/C substitution in *G8R* INR leads to a significant reduction in the 5’ poly(A) leader lengths. Bars represent the percentages of 5’ polyadenylated transcripts among all mRNAs transcribed from the *G8R^N^, G8R^P^*, and *G8R^PM^* promoters. Error bars depict 95% confidence intervals; *** indicates *p*<0.001. **(D)** Single A/C substitution in *G8R* INR increases the occurrence of 5’ capped transcripts. Bars represent the percentages of 5’ capped mRNAs transcribed from the *G8R, G8R^P^*, and *G8R^PM^* promoters. Error bars depict 95% confidence intervals; * indicates *p*<0.05. In total, 61 sequences were used for the analyses depicted in panels B-D (Tables S4, S5).

VACV was among the first DNA viruses in which 5’ capped viral mRNAs were discovered almost four decades ago, and since then, there has been a common belief that all VACV transcripts of early and postreplicative genes are capped at their 5’ ends. These findings inspired numerous fundamental investigations and discoveries on 5’ mRNA cap structures and capping machineries in other viruses and eukaryotic organisms. Our work revises this paradigm and reports a more accurate view of the transcription-coupled regulation of viral mRNA processing within different GTCs. We demonstrate here that 5’ cap occurrence in viral mRNAs gradually decreases in each successive GTC, in contrast to the reciprocal increase in 5’ poly(A) leader lengths, and that these two variables are mutually negatively correlated. We also demonstrate that the INR element directly or indirectly influences both the frequency of 5’ mRNA capping and the occurrence of 5’ poly(A) leaders, including their lengths in postreplicative VACV mRNAs. Considering all the results together, we can speculate that the degree of 5’ mRNA polyadenylation can directly affect the synthesis of the 5’ cap by some hitherto unknown mechanism. This idea is further supported by our observation that 5’ poly(A) leaders in m^7^G cap-containing VACV late transcripts were significantly shorter than the 5’ mRNA leaders, these lengths of which were calculated from the unbiased set of all VACV late mRNAs. Collectively, our results support the hypothesis that VACV transcription regulation ensures a gradual shift in viral mRNA translation initiation from a cap-dependent to cap-independent mechanism (*25, 31*), which is accompanied by virus-induced modification of the host translation machinery (*32*).

## Acknowledgments

We thank Petra Studnickova, Natalie Suchankova, Monika Kaplanova and Libor Krasny for their generous help.

## Funding

Funding was provided by the Czech Science Foundation (grant no. GA19-13491S, to T.M.); the ELIXIR CZ research infrastructure project (MEYS grant no. LM2018131, to M.P.), including access to computing and storage facilities; and the MICOBION project funded by EU H2020 (project no. 810224, to M.P.).

## Author contributions

M.S., V.V. and M.P. devised the project, main conceptual ideas and methods of data analysis; M.S., Z.M. and V.V. performed experiments; V.V. and M.S. created the figures; V.V., M.S. and M.P. analyzed data; and V.V. wrote the original draft. All authors discussed the results and edited the manuscript.

## Competing interests

The authors declare no conflicts of interest.

## Supplementary Materials

Materials and Methods

Figures S1-S6

Tables S1-S8

References (1-11)

## Notes

### Competing Interest Statement

The authors have declared no competing interest.

## References

1. G. McFadden, Poxvirus tropism. Nat Rev Microbiol 3, 201 (Mar, 2005).

2. E. Petersen et al., Human Monkeypox: Epidemiologic and Clinical Characteristics, Diagnosis, and Prevention. Infect Dis Clin North Am 33, 1027 (Dec, 2019).

3. WHO, Monkeypox - fact sheets. https://www.who.int/news-room/fact-sheets/detail/monkeypox, (2019).

4. C. R. MacIntyre, Reevaluating the Risk of Smallpox Reemergence. Military medicine, (May 6, 2020).

5. S. Melamed, T. Israely, N. Paran, Challenges and Achievements in Prevention and Treatment of Smallpox. Vaccines 6, (Jan 29, 2018).

6. S. R. Walsh, R. Dolin, Vaccinia viruses: vaccines against smallpox and vectors against infectious diseases and tumors. Expert review of vaccines 10, 1221 (Aug, 2011).

7. C. J. Baldick, Jr., B. Moss, Characterization and temporal regulation of mRNAs encoded by vaccinia virus intermediate-stage genes. J Virol 67, 3515 (Jun, 1993).

8. C. Grimm et al., Structural Basis of Poxvirus Transcription: Vaccinia RNA Polymerase Complexes. Cell 179, 1537 (Dec 12, 2019).

9. B. Y. Ahn, B. Moss, RNA polymerase-associated transcription specificity factor encoded by vaccinia virus. Proc Natl Acad Sci U S A 89, 3536 (Apr 15, 1992).

10. C. M. Wei, B. Moss, Methylated nucleotides block 5’-terminus of vaccinia virus messenger RNA. Proc Natl Acad Sci U S A 72, 318 (Jan, 1975).

11. R. F. Boone, B. Moss, Methylated 5’-terminal sequences of vaccinia virus mRNA species made in vivo at early and late times after infection. Virology 79, 67 (Jun 1, 1977).

12. O. J. Kyrieleis, J. Chang, M. de la Pena, S. Shuman, S. Cusack, Crystal structure of vaccinia virus mRNA capping enzyme provides insights into the mechanism and evolution of the capping apparatus. Structure 22, 452 (Mar 04, 2014).

13. H. S. Hillen et al., Structural Basis of Poxvirus Transcription: Transcribing and Capping Vaccinia Complexes. Cell 179, 1525 (Dec 12, 2019).

14. Z. Yang, D. P. Bruno, C. A. Martens, S. F. Porcella, B. Moss, Genome-wide analysis of the 5’ and 3’ ends of vaccinia virus early mRNAs delineates regulatory sequences of annotated and anomalous transcripts. J Virol 85, 5897 (Jun, 2011).

15. C. J. Baldick, Jr., J. G. Keck, B. Moss, Mutational analysis of the core, spacer, and initiator regions of vaccinia virus intermediate-class promoters. J Virol 66, 4710 (Aug, 1992).

16. S. S. Broyles, Vaccinia virus transcription. J Gen Virol 84, 2293 (Sep, 2003).

17. A. J. Davison, B. Moss, Structure of vaccinia virus late promoters. J Mol Biol 210, 771 (Dec 20, 1989).

18. Z. Yang et al., Expression profiling of the intermediate and late stages of poxvirus replication. J Virol 85, 98–99 (Oct, 2011).

19. C. F. Wright, B. Moss, In vitro synthesis of vaccinia virus late mRNA containing a 5’ poly(A) leader sequence. Proc Natl Acad Sci U S A 84, 8883 (Dec, 1987).

20. C. Bertholet, E. Van Meir, B. ten Heggeler-Bordier, R. Wittek, Vaccinia virus produces late mRNAs by discontinuous synthesis. Cell 50, 153 (Jul 17, 1987).

21. B. Schwer, P. Visca, J. C. Vos, H. G. Stunnenberg, Discontinuous transcription or RNA processing of vaccinia virus late messengers results in a 5’ poly(A) leader. Cell 50, 163 (Jul 17, 1987).

22. C. J. Baldick, Jr., B. Moss, Resistance of vaccinia virus to rifampicin conferred by a single nucleotide substitution near the predicted NH2 terminus of a gene encoding an Mr 62,000 polypeptide. Virology 156, 138 (Jan, 1987).

23. B. Moss, Poxvirus vectors: cytoplasmic expression of transferred genes. Curr Opin Genet Dev 3, 86 (Feb, 1993).

24. B. Schwer, H. G. Stunnenberg, Vaccinia virus late transcripts generated in vitro have a poly(A) head. Embo J 7, 1183 (Apr, 1988).

25. P. Dhungel, S. Cao, Z. Yang, The 5’-poly(A) leader of poxvirus mRNA confers a translational advantage that can be achieved in cells with impaired cap-dependent translation. PLoS pathogens 13, e1006602 (Aug, 2017).

26. M. Sýkora et al., Transcription apparatus of the yeast virus-like elements: Architecture, function, and evolutionary origin. PLoS pathogens 14, e1007377 (Oct 22, 2018).

27. V. Vopalensky, M. Sykora, T. Masek, M. Pospisek, Messenger RNAs of Yeast Virus-Like Elements Contain Non-templated 5’ Poly(A) Leaders, and Their Expression Is Independent of eIF4E and Pab1. Frontiers in microbiology 10, 2366 (2019).

28. Y. Furuichi, Discovery of m(7)G-cap in eukaryotic mRNAs. Proc Jpn Acad Ser B Phys Biol Sci 91, 394 (2015).

29. T. Urushibara, Y. Furuichi, C. Nishimura, K. Miura, A modified structure at the 5’- terminus of mRNA of vaccinia virus. FEBS Lett 49, 385 (Jan 1, 1975).

30. Z. Yang, C. A. Martens, D. P. Bruno, S. F. Porcella, B. Moss, Pervasive initiation and 3’- end formation of poxvirus postreplicative RNAs. J Biol Chem 287, 31050 (Sep 7, 2012).

31. J. Mulder, M. E. Robertson, R. A. Seamons, G. J. Belsham, Vaccinia virus protein synthesis has a low requirement for the intact translation initiation factor eIF4F, the cap-binding complex, within infected cells. J Virol 72, 8813 (Nov, 1998).

32. S. Jha et al., Trans-kingdom mimicry underlies ribosome customization by a poxvirus kinase. Nature 546, 651 (Jun 29, 2017).

33. K. Maruyama, S. Sugano, Oligo-capping: a simple method to replace the cap structure of eukaryotic mRNAs with oligoribonucleotides. Gene 138, 171 (Jan 28, 1994).

