## Supplementary Materials for "Transcripts of vaccinia virus postreplicative genes do not contain a 5’ methylguanosine cap"

**This PDF file includes:**

Materials and Methods
Figs. S1 to S6
Tables S1 to S8

### Materials and Methods

Cells

The epithelial cell lines HeLa (human cervical carcinoma) and BSC-40 (African green monkey kidney cells) were grown in Dulbecco's modified Eagle's medium (DMEM; glucose 4.5 g/l) supplemented with 10% heat-inactivated fetal bovine serum (10% FBS-DMEM) and neonatal calf serum (10% NCS-DMEM), respectively. The media contained penicillin ($1x10^5$ U/l) and streptomycin (100 mg/l). Cells were maintained at 37°C in a 5% $CO_2$ atmosphere and 95% humidity. All media and growth supplements were purchased from Gibco BRL or Sigma-Aldrich.

Viruses, infection and transfection

A purified stock of VACV strain Western Reserve (WR) was used throughout the study. The virus was propagated in BSC-40 cells supplemented with 2% NCS-DMEM as described previously (Kalbacova et al., 2008; Liskova et al., 2011) and purified by sucrose gradient sedimentation (Joklik, 1962). The titer of the virus was determined by serial dilution and plaque assays in BSC-40 cells.

For experiments, $1.5x10^6$ cells were seeded in 60-mm plates a day before infection. Cells were either mock-infected or infected at a multiplicity of infection (m.o.i.) of 5 for 40 min, washed with DMEM and supplemented with 10% FBS-DMEM (Humlova et al., 2002). When appropriate, cells were transiently transfected with the respective plasmids at 5-10 µg/plate using polyethylenimine according to (Hsu and Uludag, 2012). The culture medium was removed, and the cells were lysed in RNA Blue (Exbio) at the indicated times after infection (4 or 12 hours post infection).

RNA purification, reverse transcription and 5' RACE

Total RNA from RNA blue lysis solution was purified according to the manufacturer's protocol. DNA was removed using a DNA-free Kit (Ambion) according to the manufacturer's protocol. The integrity of the total RNA was analyzed using agarose electrophoresis as described in (Masek et al., 2005). 5' RACE-PCR experiments were performed exactly according to (Vopalensky et al., 2019). Briefly, SuperScript III reverse transcriptase, which belongs to a group of reverse transcriptases that are able to overcome the 5'–5' bond between the mRNA cap structure and the very first mRNA nucleotide (Schmidt and Mueller, 1999), was used for cDNA synthesis from total RNA using random hexamer primers (Invitrogen). After reverse transcription, the cDNA was purified using a High Pure PCR Product Purification Kit (Roche) and used for subsequent 5' RACE-PCR amplification with gene-specific primers (Table S1). Oligo-capping experiments were performed exactly according to (Vopalensky et al., 2019). The PCR amplicons were separated by agarose gel electrophoresis; each specific product was purified from the gel, inserted into the pCR4-TOPO vector (Invitrogen) and sequenced. At least 15 independent clones for individual VACV gene transcripts were analyzed. All primers used in this study are listed in Table S1.

Reporter plasmid construction

The promoter-less plasmid pEGFP-N1(-P) (Vopalensky et al., 2008) was digested with the *BamHI* and *SalI* restriction endonucleases to insert either the VACV *G8R* intermediate promoter sequence (*G8R*$^{P}$; using VV_G8Rp_1/VV_G8Rp_2 oligonucleotides) or the *G8R*$^{MP}$ promoter bearing a single point mutation within the INR (oligonucleotides VV_G8Rp_mut_1/VV_G8Rp_mut_2). Plasmids were verified by sequencing.

Statistical analyses

Statistical analyses were performed exactly according to (Vopalensky et al., 2019). Briefly, the variance in the 5' poly(A) leader lengths of selected VACV mRNAs was analyzed using the nonparametric *Kruskal-Wallis* test followed by the *post hoc Dunn* test with *p*-value adjustment according to the *Benjamini-Hochberg FDR* method. These data did not follow a normal distribution according to the *Shapiro-Wilk* test (Figs. 3; 4B; S6). Categorical binary data of 5' mRNA cap and 5' poly(A) occurrence in VACV transcripts were evaluated using two-tailed Fisher's exact test with the 95% confidence interval calculated using the adjusted Wald method (Figs. 4C, 4D).

*A5R*

114145 ATTGCGATTATAAGATTAAATG 114166

AAAAAAAAAATG
**G**AAAAAAATG
**G**AAAAAAATG
AAAAAAATG
AAAAAAATG
**G**AAAAATG
**G**AAAAATG
AAAATG
AAAATG
AAAATG
AAAATG
**G**AAATG
**G**AAATG
**G**AAATG
**G**AAATG
**G**ATG
**G**ATG

*D12L*

C 108214 ACAAACTATAGAGTTGTAAATG 108193 C

**G**AAAAAAATG
AAAAAAAATG
AAAAAAATG
AAAAAAATG
AAAAAAATG
**G**AAAAAATG
AAAAAATG
AAAAAATG
**G**AAAAATG
AAAAATG
AAAATG
**G**AAATG
**G**AAATG
**G**AAATG
AAATG

*I4L*

C 64267 AATAATTTTTTTATTACACCAA 64246 C

**G**ATTTACACCAA
**G**ATTTACACCAA
**G**ATTTACACCAA
ATTTACACCAA
ATTTACACCAA
**G**ATTACACCAA
**G**ATTACACCAA
**G**ATTACACCAA
**G**ATTACACCAA
**G**ATTACACCAA
**G**ATTACACCAA
**G**ATTACACCAA
ATTACACCAA
ATTACACCAA
ATTACACCAA
ATTACACCAA
ATTACACCAA
TTACACCAA
**G**ACACCAA

*J6R*

83286 TATTATTTTTATAGTTGTAATA 83307

**G**ATAGTTGTAATA
**G**ATAGTTGTAATA
**G**ATAGTTGTAATA
**G**ATAGTTGTAATA
**G**ATAGTTGTAATA
**G**ATAGTTGTAATA
**G**AGTTGTAATA
**G**AGTTGTAATA
**G**AGTTGTAATA
**G**AGTTGTAATA
**G**AGTTGTAATA
**G**AGTTGTAATA
**G**AGTTGTAATA
**G**AGTTGTAATA
**G**AGTTGTAATA
**G**AGTTGTAATA
**G**AGTTGTAATA
**G**AGTTGTAATA
**G**AGTTGTAATA
**G**GTTGTAATA

*H5R*

↓

91781 ACAAAGGTTCTTGAGGGTTGTG~68nts~ATG 91873

**G**GAGGGTTGTG~68nts~ATG
**G**GAGGGTTGTG~68nts~ATG
**G**GAGGGTTGTG~68nts~ATG
**G**GAGGGTTGTG~68nts~ATG
**G**GAGGGTTGTG~68nts~ATG
**G**GAGGGTTGTG~68nts~ATG
**G**GAGGGTTGTG~68nts~ATG
**G**GAGGGTTGTG~68nts~ATG
**G**GAGGGTTGTG~68nts~ATG
**G**GAGGGTTGTG~68nts~ATG
**G**GAGGGTTGTG~68nts~ATG
**G**GAGGGTTGTG~68nts~ATG
**G**GAGGGTTGTG~68nts~ATG
**G**GAGGGTTGTG~68nts~ATG
**G**GAGGGTTGTG~68nts~ATG
**G**GAGGGTTGTG~68nts~ATG
**G**GAGGGTTGTG~68nts~ATG
**G**GAGGGTTGTG~68nts~ATG
**G**GAGGGTTGTG~68nts~ATG
**G**GAGGGTTGTG~68nts~ATG

68 nts = TTAAATTGAAAGCGAGAAATAATCATAAATTATTTCATTATCGCGATATCCGTTAAGTTTGTATCGTAATG

**Fig. S1**. **5' RACE analysis of selected VACV early gene transcripts**.
The upper sequence in each panel corresponds to the viral template DNA. The TSS (black arrow) and INR (underlined) were annotated according to (Yang et al., 2011; Yang et al., 2012). If our TSS annotation differs, it is marked by an orange arrow. The sequences depicted below the template DNA represent individual sequenced cDNA clones. At least 15 independent cDNA clones were sequenced and aligned with the VACV genomic DNA sequence. Nucleotides identical to the viral template DNA are labeled in red. Nucleotides added in a nontemplated manner are labeled in green. The guanosine residues corresponding to the 5' mRNA cap are marked in black. All sequences are shown in the 5'-to-3' orientation, regardless of their transcriptional orientation in the VACV genome.

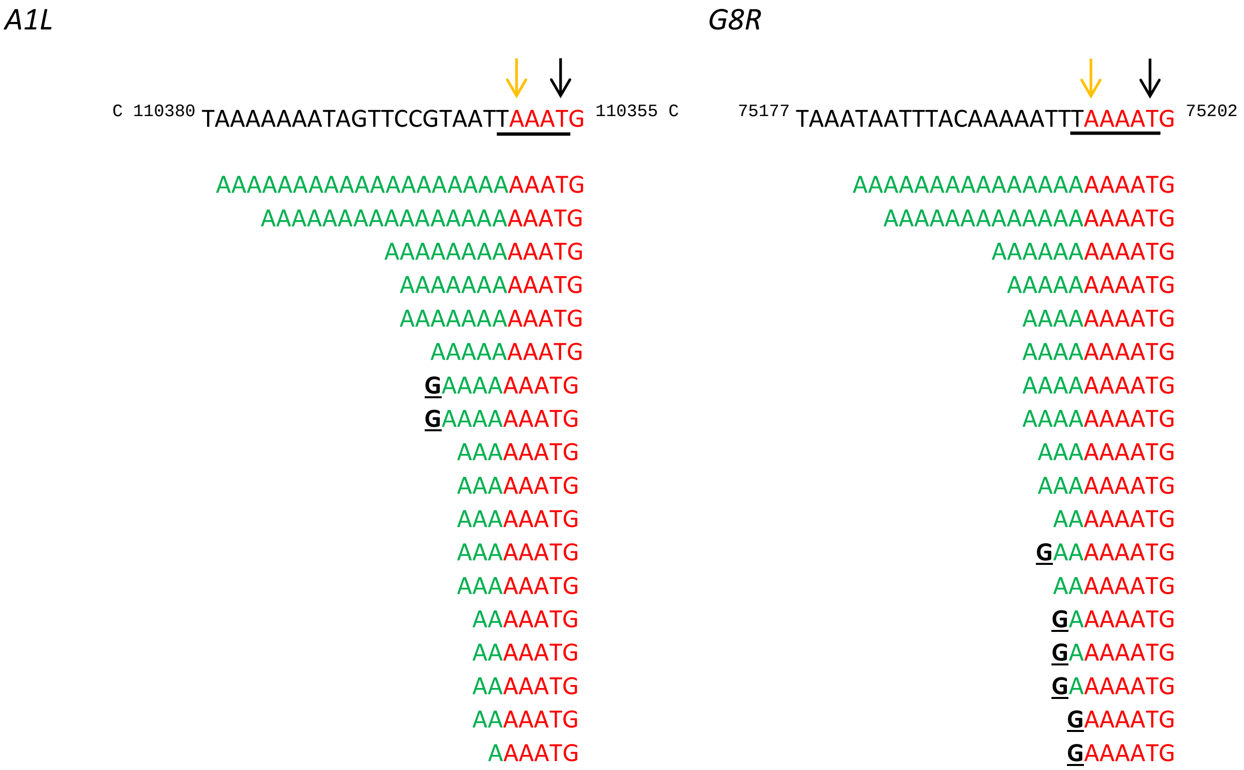


Fig. S2. **5' RACE analysis of selected VACV intermediate transcripts**.
The upper sequences correspond to the viral template DNA with the TSS annotated according to (Yang et al., 2012) (black arrow) and by us (orange arrow). The INR (underlined) is annotated according to (Yang et al., 2012). The sequences depicted below represent individual sequenced cDNA clones. The 5' untranslated regions are shown up to the ATG translation start codon. At least 18 independent cDNA clones were sequenced and aligned with the viral template. The color-coding of the figure and cDNA strand orientation are the same as those in Figure S1.

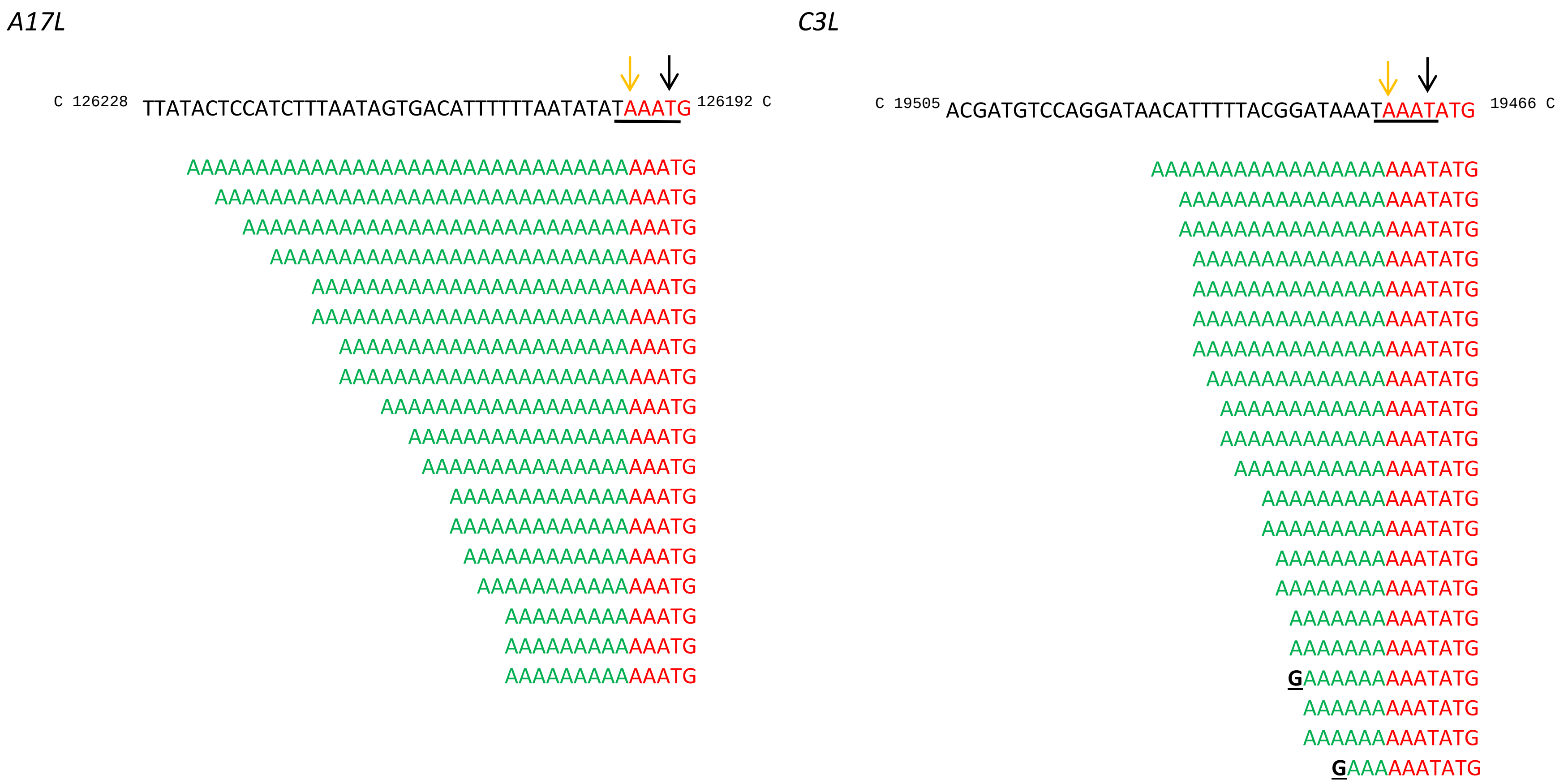

**Fig. S3. 5' RACE analysis of selected VACV late transcripts.**

The upper sequences correspond to the viral template DNA with a TSS annotated according to (Yang et al., 2012) (black arrow) and by us (orange arrow). The INR (underlined) is annotated according to (Yang et al., 2012). The sequences depicted below represent individual sequenced cDNA clones. The 5' untranslated regions are shown up to the ATG translation initiation codon. The color-coding of the figure and cDNA strand orientation are the same as those in Figure S1. At least 18 independent clones were sequenced and aligned with the viral DNA.

*A17L* (RLM-RACE)

C 126228 TTATACTCCATCTTTAATAGTGACATTTTTTAATATATAAATG 126192 C

AAAAAAAAAAAAAAAAAAAAAAAAAAAAAAAATG
AAAAAAAAAAAAAAAAAAAAAAAAAAAAATG
AAAAAAAAAAAAAAAAAAAAAAAAAAAATG
AAAAAAAAAAAAAAAAAAAAAAAAAAATG
AAAAAAAAAAAAAAAAAAAAAAAAAATG
AAAAAAAAAAAAAAAAAAAAAAAAAATG
AAAAAAAAAAAAAAAAAAAAAAAAAATG
AAAAAAAAAAAAAAAAAAAAAAAAAATG
AAAAAAAAAAAAAAAAAAAAAAAAATG
AAAAAAAAAAAAAAAAAAAAAAAAATG
AAAAAAAAAAAAAAAAAAAAAAAATG
AAAAAAAAAAAAAAAAAAAAAAAATG
AAAAAAAAAAAAAAAAAAAAAAAATG
AAAAAAAAAAAAAAAAAAAAATG
AAAAAAAAAAAAAAAAAATG
AAAAAAAAAAAAAAATG
AAAAAAAAAAAAAATG
AAAAAAAAAAAAAATG
AAAAAAAAAAAAATG
AAAAAAAAAAAAATG
AAAAAAAAAAAATG
AAAAAAAAAAAATG
AAAAAAAAAATG
AAAAAAAAAATG
AAAAAAAAATG
AAAAAAATG
AAAAAAATG
AAAAAAATG
AAAATG

*C3L* (RLM-RACE)

C 19505 ACGATGTCCAGGATAACATTTTTACGGATAAATAAATATG 19466 C

AAAAAAAAAAAAAAAAAAAAAAAATATG
AAAAAAAAAAAAAAAAAAAAAATATG
AAAAAAAAAAAAAAAAAAAAAATATG
AAAAAAAAAAAAAAAAAAAAAATATG
AAAAAAAAAAAAAAAAAAAAATATG
AAAAAAAAAAAAAAAAAAAATATG
AAAAAAAAATATG
AAAAAAAAATATG
AAAAAAAATATG
AAAAATATG
AAAAATATG
AAAAATATG
AAAAATATG
AAAAATATG
AAAAATATG
AAAAATATG
AAAAATATG
AAATATG
AAATATG
AAATATG
AAATATG
AAATATG
AAATATG
AAATATG
AAATATG
AAATATG
AAATATG
AAATATG

**Fig. S4. 5' RLM-RACE analysis of the VACV *A17L* and *C3L* late transcripts.** INRs (underlined) were annotated within the upper sequences of the viral template DNA according to (Yang et al., 2012). The sequences below represent individual sequenced cDNA clones obtained by the 5' RLM-RACE method (oligocapping). The 5' untranslated regions are shown in full up to the ATG translation start codon. At least 18 independent cDNA clones were sequenced and aligned with the viral template DNA. Sequence parts that share nucleotide identity with the template viral DNA are labeled in red, and sequence parts that are not identical to the template viral DNA are labeled in green.

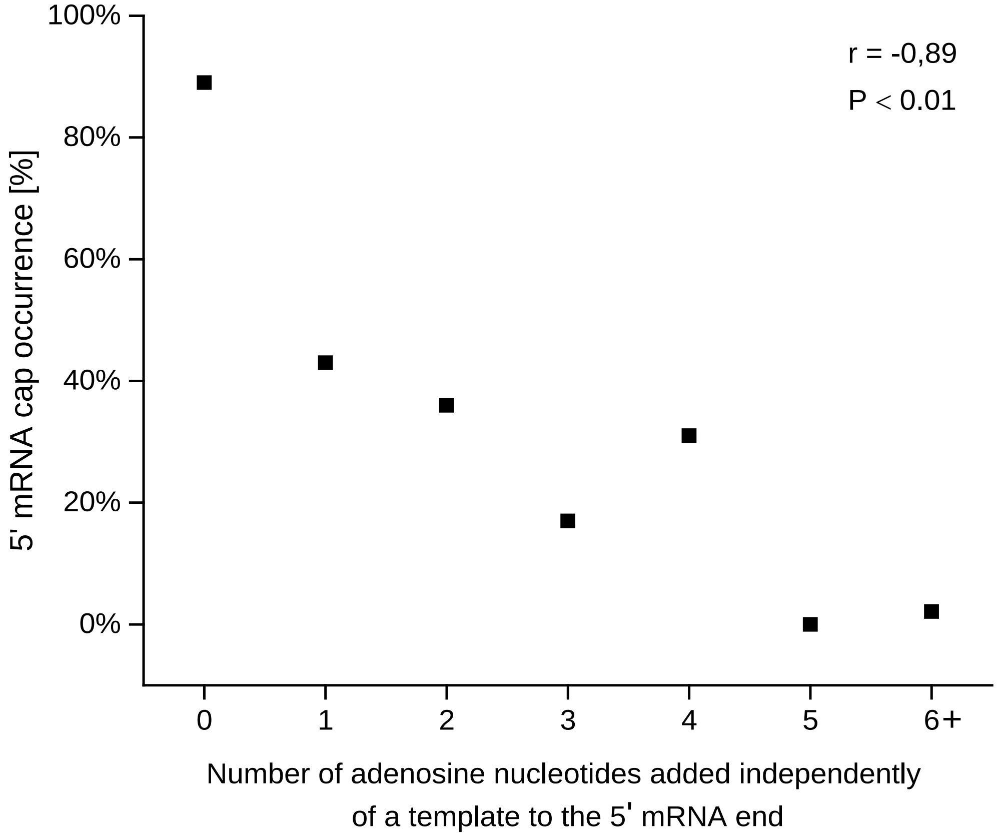


**Fig. S5. Negative correlation between the number of nontemplated adenosines in the 5' poly(A) leader and the presence of a 5' mRNA cap at the end of VACV mRNAs**. The scatter graph shows the proportion of 5' mRNA cap structures occurring in the transcripts of all GTCs. The *Pearson value correlation coefficient* is depicted as the r value. The results are significant at $p < 0.01$. In total, 166 sequences were used for this analysis (Table S6).

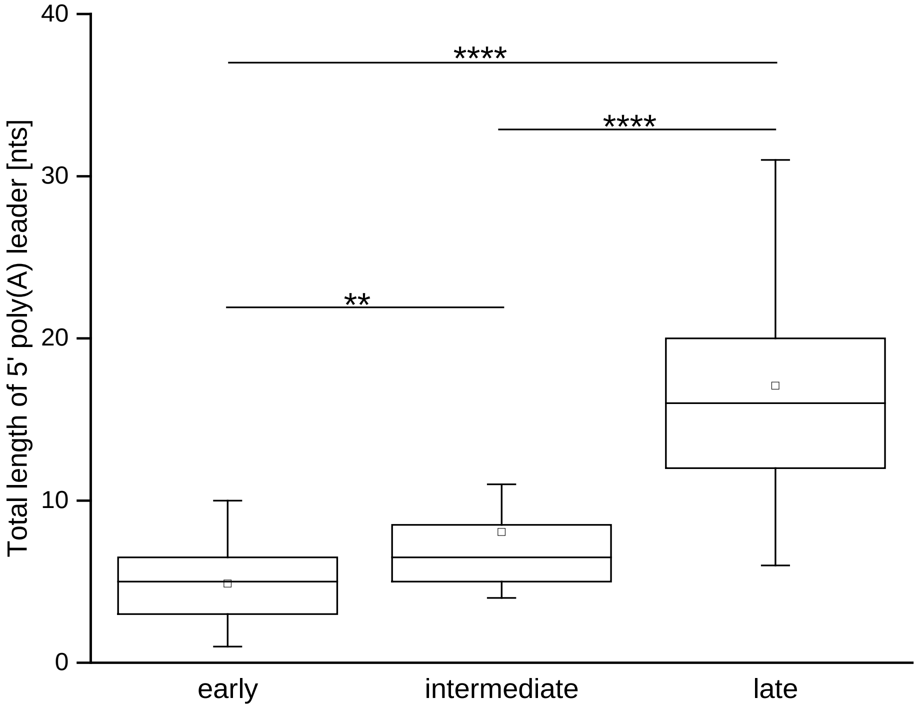


**Fig. S6 Increasing length of the 5' poly(A) leaders in mRNAs of each successive VACV GTC.** The box-whisker plot represents the total length (both templated and nontemplated adenosines) of 5' poly(A) leaders in transcripts of VACV early, intermediate and late genes. The lower bar, upper bar, bottom and top of the boxes represent the $10^{th}$ percentile, $90^{th}$ percentile and the first and third quartiles, respectively. The square dot and inner line in each box represent the mean and median, respectively. ** and **** indicate $p$ <0.01 and $p$ <0.0001, respectively. In total, 107 cDNAs corresponding to gene promoters containing the initiator region were used for this analysis (Table S6).

| Gene | Primer name | Sequence (5'-3') | Used in |
|---|---|---|---|
| A1L | 5RACE_VV_A1L | CGTAAACGCCGTCTTTATCTC | 5' RACE |
| A5R | 5RACE_VV_A5R | AGAACCCTCCTCTATCTCTTG | 5' RACE |
| A17L | 5RACE_VV_A17L | CTTCTATAGTCCTGTCTTTCG | 5' RACE / oligo-capping |
| C3L | 5RACE_VV_C3L | CCTCGGGATTCCATACCATAG | 5' RACE / oligo-capping |
| D12L | 5RACE_VV_D12L | CACGTCGAAGGTTAACATCTT | 5' RACE |
| G8R | 5RACE_VV_G8R | GAATGACGGTTCTACCACAAC | 5' RACE |
| | VV_G8Rp_1 | TCGACCATTTAACTTTAAATAATTTACAAAAATTTAAAATG | *$pG8R^{N}$*-EGFP reporter vector construction |
| | VV_G8Rp_2 | GATCCATTTTAAATTTTTGTAAATTATTTAAAGTTAAATGG | |
| $G8R^{PM}$ | VV_G8Rp_mut_1 | TCGACCATTTAACTTTAAATAATTTACAAAAATTTAACATG | *$pG8R^{PM}$*-EGFP reporter vector construction |
| | VV_G8Rp_mut_2 | GATCCATGTTAAATTTTTGTAAATTATTTAAAGTTAAATGG | |
| H5R | 5RACE_VV_H5R | TTACCAGCTTCAACTTGTACC | 5' RACE |
| I4L | 5RACE_VV_I4L | GGTCTTTCAACGATCTTGTTG | 5' RACE |
| J6R | 5RACE_VV_J6R | GGTTGCATACATTCACTGTTC | 5' RACE |
| universal | olig2(dC)anchor | GACCACGCGTATCGATGTCGACCCCCCCCCCCC | 5' RACE |
| universal | 5' RACE Adapter | GCUGAUGGCGAUGAAUGAACACUGCGUUUGCUGGCUUUGAUGAAA | oligo-capping |
| universal | 5' RACE Outer Primer | GCTGATGGCGATGAATGAACACTG | oligo-capping |
| universal | 5' RACE Inner Primer | CGCGGATCCGAACACTGCGTTTGCTGGCTTTGATG | oligo-capping |

**Table S1. Primers used in this study.**

| Vaccinia gene time class | Gene | IN R | Number of 5' nontemplated adenosines | Minimal number of adenosines[1] at 5' ends of uncapped transcripts | mRNAs with 5' nontemplated poly(A) leaders [%] | Median/Mean of added adenosines per mRNA molecule | 5' capped mRNAs [%] | Number of analyzed clones |
|---|---|---|---|---|---|---|---|---|
| Early | ***J6R*** | no | 0 | 0 | 0 | 0/0 | 100 | 20 |
| | ***H5R*** | no | 0 | 0 | 0 | 0/0 | 100 | 20 |
| | ***I4L*** | no | 0–1 | 1 | 63.1 | 0/0.3 | 52.6 | 19 |
| | ***A5R*** | yes | 0–7 | 4 | 64.7 | 1/1.7 | 58.8 | 17 |
| | ***D12L*** | yes | 0–5 | 3 | 73.3 | 3/2.3 | 40.0 | 15 |
| Intermediate | ***G8R*** | yes | 6–15 | 6 | 88.9 | 3/3.9 | 33.3 | 18 |
| | ***A1L*** | yes | 1–19 | 4 | 100 | 3/5.2 | 11.1 | 18 |
| Late | ***C3L*** | yes | 9–17 | 9 | 100 | 11/10.5 | 9.5 | 21 |
| | ***A17L*** | yes | 9–32 | 9 | 100 | 17/18.3 | 0 | 18 |

[1] templated and nontemplated

**Table S2. Summary of the 5' RACE experiments.** Each VACV gene is characterized by the gene name; presence or absence of the INR; number of adenosines added in a nontemplated manner; minimal number of total adenosines (both templated and nontemplated) at the 5' ends of uncapped transcripts; percentage of all transcripts containing 5' nontemplated adenosine nucleotides; the average number (shown as the median and mean) of nontemplated adenosine nucleotides per mRNA molecule; percentage of all transcripts containing a 5' cap; and number of analyzed cDNA clones. Related to Figures 1 and 2.

| Vaccinia gene time class | Method | Gene | Number of 5' nontemplated adenosines | mRNAs with 5' nontemplated poly(A) leaders [%] | Median/Mean of added adenosines per mRNA molecule | 5' capped mRNAs [%] | Number of analyzed clones |
|---|---|---|---|---|---|---|---|
| **Late** | **5' RACE** | *C3L** | 9–17 | 100 | 11/10.5 | 9.5 | 21 |
| | | *A17*L* | 9–32 | 100 | 17/18.3 | 0 | 18 |
| **Late** | **RLM-RACE** | *C3L* | 0–21 | 60.7 | 2/5.2 | N/A | 28 |
| | | *A17L* | 1–28 | 100 | 14/14.2 | N/A | 29 |

[1] templated and nontemplated
* data from Table S2

**Table S3**. **5' RLM-RACE analysis of VACV *A17L* and *C3L* late transcripts**. The results of 5' RLM-RACE in combination with the results of classic 5' RACE-PCR (data from Table S2) show that transcripts of both selected VACV late genes containing the 5' mRNA cap structure have shorter 5' poly(A) leaders than uncapped transcripts of the same genes. Related to Figure 3.

| Vaccinia gene time class | Promoter | Number of 5' nontemplated adenosines | Minimal number of adenosines[1] at 5' end of uncapped transcripts | mRNAs with 5' nontemplated poly(A) leaders [%] | Median/Mean of added adenosines per mRNA molecule | 5' capped mRNAs [%] | Number of analyzed clones |
|---|---|---|---|---|---|---|---|
| **Intermediate** | ***G8R*[N]*** | 6–15 | 6 | 88.9 | 3/3.9 | 33.3 | 18 |
| | ***G8R*[P]** | 0–9 | 3 | 83.3 | 2.5/3 | 20.8 | 24 |
| | ***G8R*[PM]** | 0–1 | 2 | 31.6 | 0/0.3 | 57.9 | 19 |

[1] templated and nontemplated
[N]* data from Table S2, native *G8R* promoter, VACV mRNA purified from infected HeLa cells
[P] plasmid-localized reporter gene (*EGFP*) under the control of the *G8R* VACV intermediate promoter
[PM] plasmid-localized reporter gene (*EGFP*) under the control of the *G8R* VACV intermediate promoter containing a single point mutation in the INR

**Table S4. VACV INR controls 5' end formation of viral mRNAs.** The results show that a single A/C substitution within the short VACV INR leads to a significant reduction in mRNA 5' poly(A) leader length and to the switching of mRNA ends from 5' polyadenylated to 5' capped. Related to Figure 4.

| | *G8R*$^{N}$* | *G8R*$^{P}$ |
|---|---|---|
| *G8R*$^{P}$ | 0.764956 | |
| *G8R*$^{PM}$ | 0.000037 | 0.000037 |

* native *G8R* promoter, VACV mRNA purified from infected HeLa cells
$^{P}$ plasmid localized reporter gene (*EGFP*) under the control of a *G8R* VACV intermediate promoter
$^{PM}$ plasmid localized reporter gene (*EGFP*) under the control of the *G8R* VACV intermediate promoter containing a single point mutation in the INR

**Table S5. The results of the statistical analysis are depicted in** Figure 4. *Dunn p*-values, further adjusted by the *Benjamini-Hochberg FDR* method, are depicted in the table above.

| Number of nontemplated adenosines | Number of sequences | Number of sequences with $m^7G$ structure | 5' mRNA cap occurrence [%] |
|---|---|---|---|
| **0** | 66 | 59 | 89 |
| **1** | 14 | 6 | 43 |
| **2** | 11 | 4 | 36 |
| **3** | 12 | 2 | 17 |
| **4** | 13 | 4 | 31 |
| **5** | 3 | 0 | 0 |
| **6+** | 47 | 1 | 2.1 |

**Table S6.** Proportion of mRNAs containing 5' cap structures among VACV mRNAs with different numbers of nontemplated adenosines in their 5' poly(A) leaders. Related to Figure S5.

| Gene time class | Number of clones | 5' mRNA cap occurrence [%] | Median/Mean of total length of 5' poly(A) leader [nts] |
|---|---|---|---|
| **early w/o INR** | 59 | 86 | -/- |
| **early with INR** | 32 | 50 | 5/5 |
| **intermediate** | 36 | 22 | 7/8 |
| **late** | 39 | 5 | 16/17 |

**Table S7.** Occurrence of 5' mRNA cap structures and length of the 5' poly(A) leader in VACV transcripts from different GTCs. Related to Figure S6.

| | **early** | **intermediate** |
|---|---|---|
| **intermediate** | 4.625424e-03 | |
| **late** | 9.461407e-15 | 3.706485e-07 |

**Table S8**. **The results of the statistical analysis are depicted in Figure S6.** *Dunn p*-values, further adjusted by the *Benjamini-Hochberg FDR* method, are depicted in the table above.

**Supplementary references**

Hsu, C.Y., and Uludag, H. (2012). A simple and rapid nonviral approach to efficiently transfect primary tissue-derived cells using polyethylenimine. Nature protocols *7*, 935-945.
Humlova, Z., Vokurka, M., Esteban, M., and Melkova, Z. (2002). Vaccinia virus induces apoptosis of infected macrophages. J Gen Virol *83*, 2821-2832.
Joklik, W.K. (1962). The purification fo four strains of poxvirus. Virology *18*, 9-18.
Kalbacova, M., Spisakova, M., Liskova, J., and Melkova, Z. (2008). Lytic infection with vaccinia virus activates caspases in a Bcl-2-inhibitable manner. Virus Res *135*, 53-63.
Liskova, J., Knitlova, J., Honner, R., and Melkova, Z. (2011). Apoptosis and necrosis in vaccinia virus-infected HeLa G and BSC-40 cells. Virus Res *160*, 40-50.
Masek, T., Vopalensky, V., Suchomelova, P., and Pospisek, M. (2005). Denaturing RNA electrophoresis in TAE agarose gels. Anal Biochem *336*, 46-50.
Schmidt, W.M., and Mueller, M.W. (1999). CapSelect: a highly sensitive method for 5' CAP-dependent enrichment of full-length cDNA in PCR-mediated analysis of mRNAs. Nucleic Acids Res *27*, e31.
Vopalensky, V., Masek, T., Horvath, O., Vicenova, B., Mokrejs, M., and Pospisek, M. (2008). Firefly luciferase gene contains a cryptic promoter. RNA *14*, 1720-1729.
Vopalensky, V., Sykora, M., Masek, T., and Pospisek, M. (2019). Messenger RNAs of Yeast Virus-Like Elements Contain Non-templated 5' Poly(A) Leaders, and Their Expression Is Independent of eIF4E and Pab1. Frontiers in microbiology *10*, 2366.
Yang, Z., Bruno, D.P., Martens, C.A., Porcella, S.F., and Moss, B. (2011). Genome-wide analysis of the 5' and 3' ends of vaccinia virus early mRNAs delineates regulatory sequences of annotated and anomalous transcripts. J Virol *85*, 5897-5909.
Yang, Z., Martens, C.A., Bruno, D.P., Porcella, S.F., and Moss, B. (2012). Pervasive initiation and 3'-end formation of poxvirus postreplicative RNAs. J Biol Chem *287*, 31050-31060.
